# Nanobubble-Based Ultrasound Localization Microscopy through Interactive Adaptive Processing

**DOI:** 10.64898/2026.07.14.738435

**Authors:** Grigori Shapiro, Yarin Gershman, Mike Bismuth, Tali Ilovitsh

## Abstract

This study presents the use of sub-micron nanobubbles (NBs) as contrast agents for ultrasound localization microscopy (ULM), a super-resolution imaging technique that visualizes microvascular structure and flow beyond the acoustic diffraction limit. While ULM has traditionally relied on micron-sized microbubbles (MBs), the reduced dimensions and prolonged circulation times of NBs make them attractive candidates for localization-based imaging. However, their weaker acoustic responses present significant challenges for reliable detection and tracking. To address this challenge, we developed the ULM Master GUI, an interactive framework for optimization of the complete ULM processing pipeline. Using custom ultrasound-compatible wall-less gelatin flow phantoms containing vessel-mimicking channels and bifurcations ranging from 100 to 500 μm, we demonstrate that NB-based ULM achieves velocity reconstruction and flow partitioning measurements comparable to conventional MB-based ULM. Across all investigated geometries, NBs faithfully reproduced the underlying flow patterns and hemodynamic behavior despite their substantially reduced acoustic scattering. These findings establish the feasibility of NB-based ULM, expand the range of contrast agents available for localization microscopy, and provide a foundation for future super-resolution ultrasound imaging using nanoscale acoustic contrast agents. The ULM processing GUI is publicly available at https://github.com/grisha1998/ulm-super-resolution-toolbox.

## Introduction

The visualization of the microvascular network is critical for the early detection and characterization of pathologies ranging from tumor neoangiogenesis to neurodegenerative diseases [1]. Conventional medical ultrasound, however, is fundamentally constrained by the acoustic diffraction limit, which dictates that spatial resolution is restricted to approximately half the acoustic wavelength [2]. This constraint creates an inherent trade-off between imaging depth and resolution, leaving deep-seated microvasculature, such as the human cerebral or abdominal microcirculation, largely inaccessible to standard B-mode or Power Doppler techniques [3]. ULM emerged as a paradigm shift to bypass this barrier, inspired by optical super-resolution techniques such as PALM and STORM [4]. This approach utilizes MBs, which are gas-filled, shell-encapsulated contrast agents typically 1.5-10 μm in diameter that serve as highly echogenic intravascular tracers [5]. In ULM, individual MB signals are isolated and localized across thousands of ultrafast imaging frames, enabling reconstruction of super-resolved vascular maps with spatial resolution far beyond the system point spread function (PSF) [6]. However, due to their micron-scale size, MBs may provide less efficient coverage of the smallest microvascular compartments, particularly capillary beds with diameters of 2-6 μm [7]. In ULM studies of the brain microvasculature, reconstruction of capillary networks requires substantially longer acquisition times than larger vessels, owing to sparse MB passages and slower blood flow in these regions [8]. Similarly, focused ultrasound-mediated blood brain barrier opening studies have reported reduced effects in the smallest capillary segments [9], [10], highlighting the challenges of achieving uniform MB-mediated bioeffects throughout the microvascular network. Furthermore, because MBs remain largely intravascular, their penetration into the extravascular tumor space is inherently limited [11], [12].

To overcome these limitations, sub-micron NBs have emerged as a promising next frontier in contrast-enhanced ultrasound. Beyond their ability to traverse narrower channels, NBs exhibit significantly longer circulation half-lives and enhanced mechanical stability compared to MBs [10], [13]. Their small size also enables improved interaction with capillary-scale vasculature and offers the potential for molecular targeting and extravasation through the leaky tumor microenvironment via the enhanced permeability and retention effect [14], [15]. However, the small size of NBs also makes them substantially more challenging to detect with ultrasound, as their weak acoustic echoes are frequently obscured by tissue clutter and electronic noise [5], [16]. This presents a key challenge for NB-based ULM. Here, we introduce NB-based ULM and demonstrate, to the best of our knowledge, the first implementation of ULM using NBs as the primary contrast agent. While NBs have been extensively investigated as contrast agents for diffraction-limited contrast-enhanced ultrasound, including molecular imaging of tumors and characterization of tissue perfusion kinetics [17], [18], [19], these studies have not pursued sub-pixel localization or trajectory-based super-resolution velocity mapping. Similarly, nanodroplet-based super-resolution approaches have demonstrated localization microscopy using phase-change agents [20], [21], however, acoustic vaporization transforms the NDs into MB-like contrast agents during imaging. To address the processing challenges associated with the weak acoustic signals of NBs, we developed the ULM Master GUI, an interactive framework that integrates the complete ULM pipeline and provides real-time visual and quantitative feedback for systematic optimization of the ULM processing pipeline. Using custom ultrasound-compatible wall-less blood vessel flow phantoms [22], [23], we then evaluated the feasibility of NB-based ULM and compared its performance with conventional MB-based ULM. These findings establish the feasibility of NB-based ULM and provide a foundation for future super-resolution imaging applications using nanoscale ultrasound contrast agents.

## Materials and Methods

### Contrast Agent Synthesis and Characterization

Lipid-shelled MBs were synthesized using a custom phospholipid formulation comprising DSPC and DSPE-PEG2K dissolved in a propylene glycol / glycerol mixture at a 1:3 ratio (high-glycerol content). The lipid phases were combined with PBS and sonicated at 62°C to ensure homogeneity, then aliquoted into 3 mL headspace vials and saturated with perfluorobutane C_4_F_10_ gas [11]. This high-glycerol formulation promotes a flexible lipid arrangement that enhances the nonlinear acoustic signature [24]. Prior to use, vials were activated by mechanical shaking (45 s, VialMix) and purified via centrifugation to isolate the target size fraction and remove excess lipid debris [25]. NBs were formulated from a four-component lipid shell, C22, DPPA, DPPE, and DSPE-PEG2K, dissolved in propylene glycol, glycerol and PBS at 80°C, owing to the elevated phase transition temperature of C22. Aliquots were saturated with Octafluoropropane C_3_F_8_ gas. Prior to use, vials were activated by mechanical shaking (45 s, VialMix). The sub-micron NB fraction was isolated from residual larger bubbles by buoyancy-based fractionation: inverted centrifugation at 500 rpm for 5 min, with the NB-enriched fraction extracted from the bottom of the inverted vial [10], [16], [26]. Post-synthesis characterization (Accusizer FXNano, Particle Sizing Systems, Entegris, MA) confirmed a mean MB diameter of 1.3±0.3 μm at a total number concentration of 1.40×10^10^ mL^−1^ (Figure 1 a) and a mean NB diameter of 168±5.5 nm at a total number concentration of 1.64×10^12^ mL^−1^ (Figure 1 b), establishing the two independent contrast agent populations required for comparative analysis.

**Figure 1:**
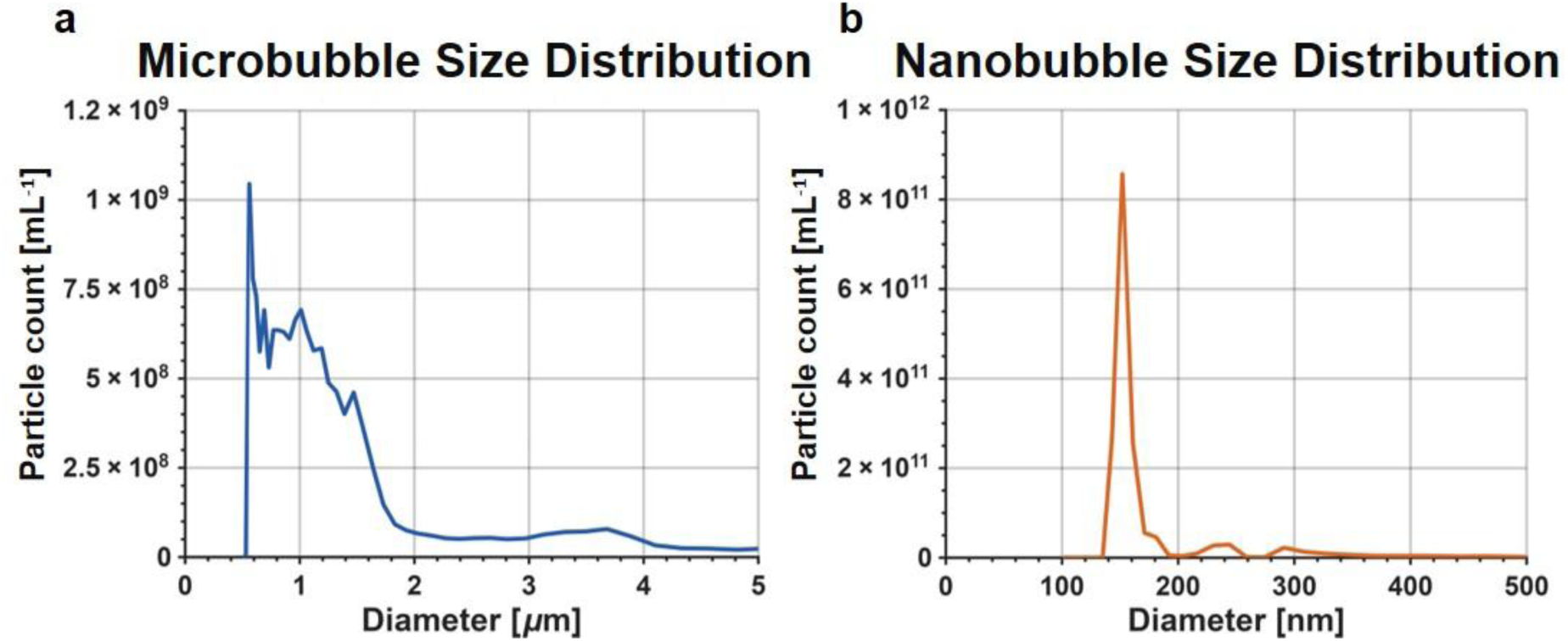
Size distribution of MB and NB. (**a**) Microbubbles. **(b)** Nanobubbles. measured by Accusizer FXNano, verifying the distinct size populations.

According to the Rayleigh scattering theory, the scattering cross-section, σ*_s_*, of a bubble much smaller than the ultrasound wavelength is proportional to the sixth power of its radius, *r* [27]:

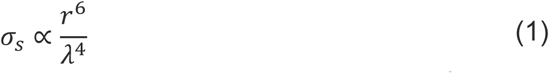

The resulting 10-fold reduction in mean radius from MBs to NBs leads to a 10^6^ decrease in scattering power, implying a theoretical backscattered intensity penalty of ∼60 dB [5], [17]. This severe signal loss motivates the development of the specialized, interactive processing framework described in this study to preserve and resolve these faint acoustic signatures [7].

### Tissue-mimicking Phantom Fabrication and Experimental Setup

Custom ultrasound-compatible wall-less blood vessel flow phantoms were fabricated as previously described in [22], [23]. The phantoms contained blood-vessel-mimicking channel geometries and served as controlled flow environments for ULM experiments. A 12% (w/v) gelatin solution (18 g in 150 mL deionized water) was selected as the matrix material for its ability to approximate the speed of sound (c ≈ 1540 m/s) and acoustic attenuation of soft tissue [22], [28], [29]. Custom wall-less flow phantoms containing vessel-mimicking channels with diameters ranging from 100 to 500 μm were fabricated using a two-part CNC-machined aluminum mold (Figure 2 a,c). The liquid gelatin was poured into the mold, gelled at 4°C for 2 h, and subsequently bonded for 24 h to form a sealed flow phantom (Figure 2 b,d). The wall-less architecture eliminated acoustic reflections from tubing interfaces, thereby minimizing interference with low-amplitude NB signals. Diluted NB or MB solutions were infused through the phantom via a programmable syringe pump (GenieTouch, Kent Scientific, Torrington) at constant flow rates between 0.01 and 0.25 mL min^−1^ [22], [23]. Both agent types were prepared via serial dilution of the post-activation stock: 5 µL were added to 10 mL PBS (×2,000), 1 mL of this intermediate was transferred to 10 mL PBS (×10), and 2 mL of the resulting solution were transferred to the final 2.5 mL syringe (×1.25), yielding a total dilution factor of 1:25,000. The solutions were utilized for experiments within 3 h of activation.

**Figure 2:**
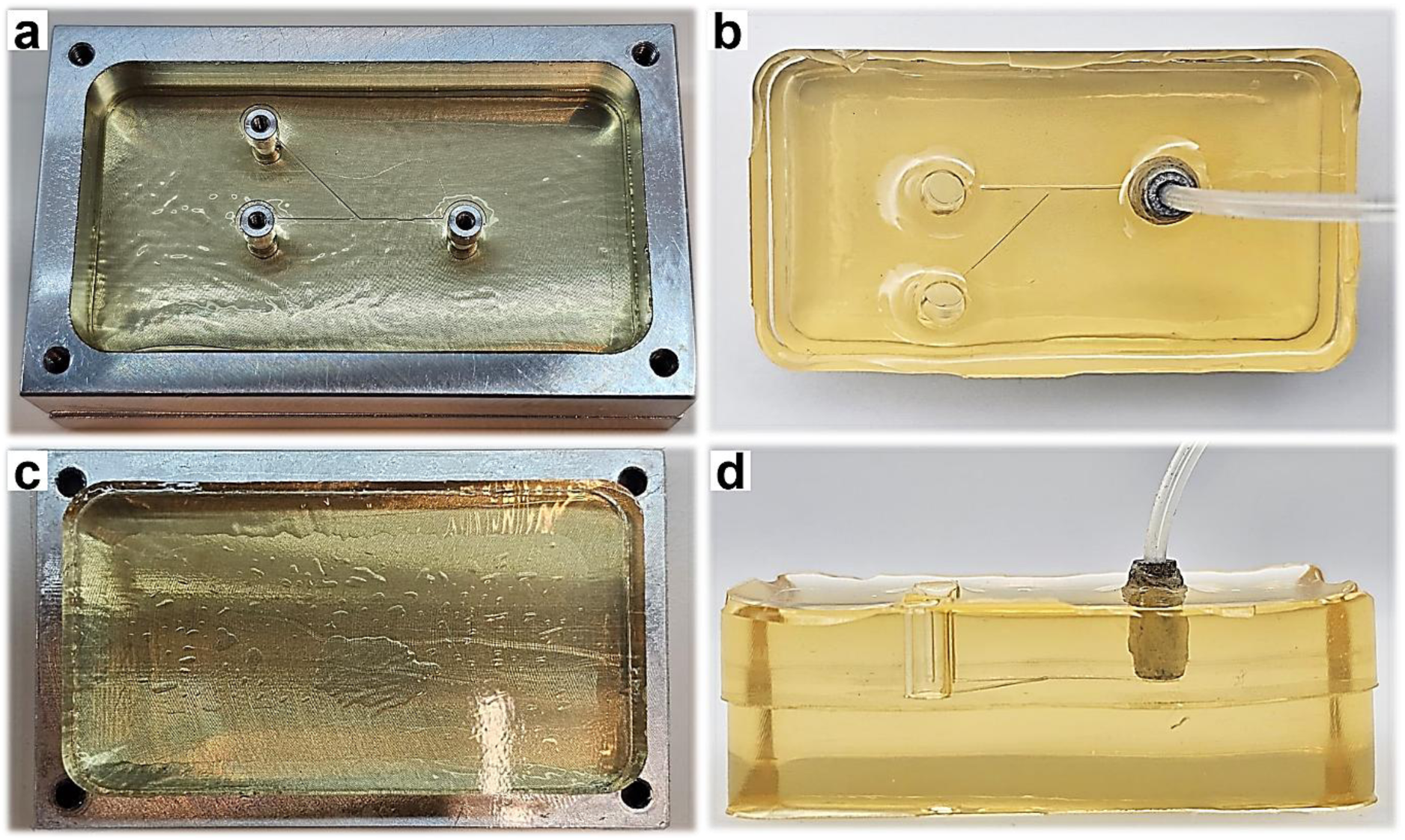
Design and assembly of the custom-made gelatin phantom. (**a**) Top mold component featuring the microvascular-mimicking channel structure and inlet/outlet connectors. **(b)** Superior view of the finalized phantom with internal vascular channels visible through the gelatin and microfluidic tubing connected for flow. **(c)** Bottom gelatin layer providing the flat bonding surface. **(d)** Lateral view of the complete dual-layer assembly.

To establish a ground truth for the flow validation, the theoretical peak flow velocity (*V_peak_*) within the phantom channels was calculated assuming a fully developed laminar flow profile (Eq. 2) [23]. In such a profile, the peak velocity at the center of the channel is approximately twice the average velocity (*V_avg_*), which is determined by the relationship between the volumetric flow rate (*Q*) and the channel’s cross-sectional area (*A*):

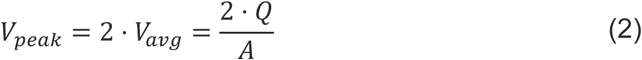

where *Q* is the volumetric flow rate programmed into the syringe pump. Since the phantom channels possess a rectangular cross-section, the area *A* was defined as *A* = *W* ⋅ *H*, where *H* is the constant channel height 300 μm and *W* is the specified channel width. These calculated values served as the baseline to evaluate the accuracy of the velocities reconstructed by the ULM processing pipeline.

### Ultrasound Data Acquisition

All US data were acquired using the Verasonics Vantage 256 programmable research system (Verasonics Inc., Kirkland, WA, USA) with the L22-14vX high-frequency linear array transducer (128 elements, 0.10 mm pitch, 18 MHz center frequency, fixed elevation focus at 8 mm). A high-frame-rate coherent plane-wave (PW) compounding sequence across three steered angles (−5°, 0°, +5°) was employed to balance spatial resolution, SNR, and temporal fidelity. Compounded IQ frames were streamed in real time into a pre-allocated RAM buffer and batch-transferred to SSD upon sequence completion, ensuring temporal continuity of bubble trajectories. The inter-frame bubble displacement was maintained at approximately 1– 2 pixels throughout all experiments. The axial imaging window was set between 14 and 20 mm depth (6 mm active zone), yielding a native pixel size of ∼49 μm in both lateral and axial dimensions. Imaging was performed at two transmit voltage levels, 20 V and 30 V. Acoustic calibration was carried out using a needle hydrophone (NH0200, Precision Acoustics, U.K.) in a degassed water tank. The lower voltage (20 V) was used for the localization algorithm comparison experiments, while both voltage levels were employed for velocity profile reconstruction and flow validation studies to evaluate NB detection sensitivity across acoustic conditions.

### ULM Processing Pipeline

ULM processing was performed using a custom MATLAB framework comprising clutter filtering, localization, tracking, and rendering stages [4], [6], [30]. Tissue clutter was removed using SVD-based filtering [31], with thresholds selected using the Spatial Similarity Matrix method [32]. Bubble locations were identified through regional-maxima detection and localized at sub-pixel resolution using either Radial Symmetry or 2D Gaussian Fitting [33], [34]. Quality-control filters were applied to reject low-confidence localizations [35]. The tracking engine supports four algorithms evaluated comparatively: Nearest Neighbor (NN, greedy baseline) [6], Global Hungarian Tracker (HT, linear assignment with the Munkres solver) [36], Kalman Tracker (KT, constant-velocity predictive state estimation with gap-closing) [35]. All trackers beyond NN can operate with a Smart Cost Matrix (SC) that augments the base spatial cost with multiplicative penalty terms. Trajectory direction is estimated via linear regression over the last *N* Kalman-smoothed positions to mitigate sub-pixel jitter, informing both a hard angular gate and a ramped directional penalty (*P_angle_*) for deviations exceeding a specified threshold. Additionally, a brightness consistency penalty (*P_intensity_*) scales with the normalized difference between the candidate intensity and the track’s incrementally cached mean brightness. This formulation substantially reduces false links at bifurcations without relying on proximity alone [37]. Reconstructed trajectories are projected onto a super-resolution grid upsampled 10× relative to native pixel size. Prior to projection, tracks are smoothed with a Savitzky-Golay filter and densely interpolated using cubic splines to suppress localization jitter while preserving vessel geometry [6]. The rendering engine produces four outputs: a vessel density map (power-law compression, *γ* = 0.3), an HSV velocity-density fusion (hue encodes velocity; brightness encodes density), and raw and lightly smoothed velocity maps [22], [23]. Rendering is decoupled from tracking, enabling rapid iterative post-processing without recomputing the full pipeline

### Interactive Optimization: The ULM Master GUI

The ULM Master GUI is a MATLAB-based interactive platform for optimization and visualization of the ULM processing pipeline (Figure 3). The GUI integrates clutter filtering, localization, tracking, and rendering within a single environment and provides visual and quantitative feedback at each processing stage. Real-time SVD filtering is implemented together with Spatial Similarity Matrix visualization [32], facilitating selection of the clutter-filtering rank range. Detected candidate events are displayed as overlays on filtered B-mode images, and localization statistics are provided for assessment of detection performance.

**Figure 3:**
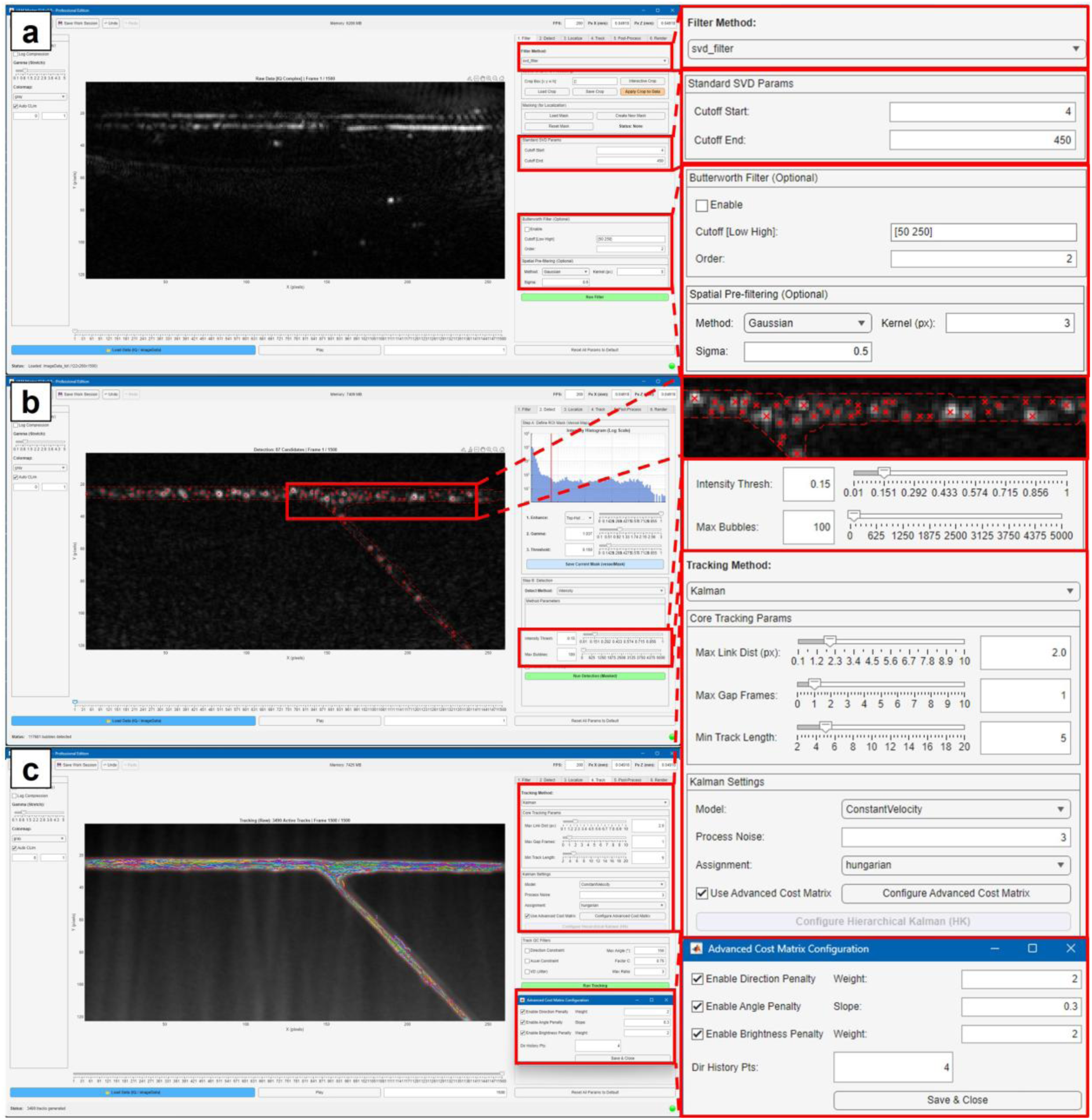
The ULM Master GUI – an interactive end-to-end optimization platform for ULM. The interface exposes six sequential processing stages (tab bar, top right) with real-time visual feedback. **(a)** Stage 1 – Clutter Filtering. Raw IQ data (Frame 1/1500) is shown with the SVD filter control panel (insets): singular value cutoff range, optional Butterworth temporal bandpass, and Gaussian spatial kernel. **(b)** Stage 2 – Detection. NB candidate localizations (red markers, inset) overlaid on the filtered B-mode frame. The dashed red boundary defines the vessel ROI mask. Detection threshold and maximum bubble count are set via the control panel (inset). **(c)** Stage 4 – Tracking. Color-encoded NB trajectories accumulated over 1500 frames on a TMIP background. The Kalman tracker uses a constant-velocity model with Hungarian assignment. The Advanced Cost Matrix Configuration panel (inset) shows the active penalty terms: Direction Penalty and Angle Penalty, which together define the soft ramped cost for trajectory deviations, and Brightness Penalty which penalizes acoustic intensity mismatches. The panel also allows for tuning the Dir History Pts, defining the number of previous smoothed positions used for the linear regression direction estimate.

Trajectories are visualized on a TMIP background to facilitate evaluation of tracking continuity, and rendering parameters can be adjusted with immediate visualization of the resulting super-resolution maps. A metadata-driven initialization step extracts acquisition parameters and automatically computes spatial calibration factors used for velocity estimation and image rendering. Optimization is performed through four sequential stages: (1) SVD rank selection using the Spatial Similarity Matrix; (2) detection-threshold adjustment using localization overlays on filtered B-mode images; (3) tracking-parameter tuning using trajectory visualization on the TMIP background; and (4) rendering-threshold adjustment on the super-resolution grid. The sequential workflow allows verification of processing outputs at each stage before proceeding to subsequent steps. The complete processing framework, including the modular ULM pipeline and interactive GUI, is implemented in MATLAB and is publicly available as an open-source toolbox (see Code Availability).

### Quantitative Performance Metrics

Four metrics were used to evaluate and compare NB and MB performance. The Tortuosity Index *τ* (Eq. 3), is defined as the ratio of actual path length to the straight-line Euclidean distance between track endpoints [38], [39], where *τ* ≈ 1 indicates smooth laminar trajectories and *τ* ≫ 1 indicates erratic, jitter-dominated paths.

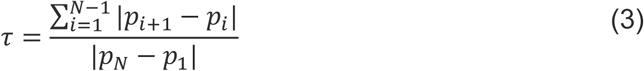

where *p_i_* represents the spatial coordinates of the localized bubble at frame *i*, and *N* is the total number of frames in the track. Track length distributions were analyzed as an indicator of temporal continuity, with a shift toward longer tracks reflecting successful gap-closing [6], [38]. Cross-sectional velocity profiles were extracted at defined phantom ROIs (pre-bifurcation, post-bifurcation, and branch segments) [23] and compared against the expected parabolic profile as a direct validation of flow-tracing fidelity. Finally, vascular partitioning was quantified via the partitioning ratio *P_branc_*_ℎ_ (Eq. 4), calculated from localization counts in three ROIs at each bifurcation. This concentration-normalized metric isolates hemodynamic behavior from the ∼100× difference in native NB versus MB dose [10], enabling a direct functional comparison of the two agents [13], [40].

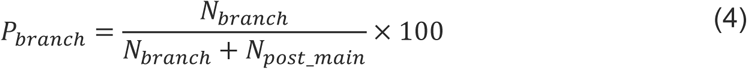

where *P_branc_*_ℎ_ represents the percentage of contrast agents diverted into the secondary branch, *N_branc_*_ℎ_ is the number of localizations detected within the branch ROI, and *N_post_*__*main*_is the number of localizations detected in the continuing main channel ROI.

## Results

### Tracking Algorithm Evaluation on NB

Following the development of the processing framework, a controlled in silico experiment was conducted to determine the optimal tracking strategy for sub-wavelength NBs. The primary objective was to evaluate how different tracking logics handle the reconstruction of complex micro-channel geometries under controlled flow conditions. To this end, lipid-shell NBs were infused into a custom ultrasound-compatible wall-less blood vessel flow phantom featuring a 300 μm primary channel that bifurcates into a 100 μm secondary branch. A syringe pump maintained a steady flow rate of 0.03 mL min^−1^, with imaging at MI = 0.32, establishing a laminar flow profile with a theoretical centerline velocity of approximately 11 mm·s^−1^. To ensure a valid comparison, all datasets were processed using identical clutter filtering and localization parameters, with a strict minimum track length filter of 15 frames applied to isolate only robust trajectories.

The impact of the tracking algorithm on the final super-resolution image is evaluated by comparing the velocity maps generated by five progressively advanced tracking configurations: Nearest Neighbor (NN), standard Hungarian Tracking (HT), standard Kalman Tracking (KT), HT with a smart cost matrix (HT SC), and KT with a smart cost matrix (KT SC) (Figure 4). The limitations of basic tracking algorithms are evident in the reconstructions using the NN (Figure 4 b-e) and the standard HT (Figure 4 f-i) methods. While both capture the basic boundary layer effect, they are limited in resolving the distinct centerline peak characteristic of laminar flow. Instead, the central flow regions appear homogeneous and noisy, exhibiting frequent velocity fluctuations that lack spatial coherence (Figure 4 b-i). Although the standard HT method improves global track continuity compared to NN, the velocity profile remains blunted, indicating an inability to resolve fine gradients. Configurations incorporating predictive motion modeling or advanced cost constraints provide visible improvements in flow visualization. The standard KT introduces a predictive motion model that smooths velocity estimation, thereby revealing the parabolic nature of the flow (Figure 4 j-m). However, tracking noise and erroneous track associations persist outside the channel boundaries in this configuration. Conversely, the HT method with a SC utilizes directional and brightness penalties, implemented within the advanced cost calculation engine, to delineate the laminar profile more clearly. Despite this clarity, the strict rejection criteria inherent to this configuration result in a sparser map, particularly within the bifurcation zone where flow directions are more complex (Figure 4 n-q). Ultimately, the optimal reconstruction is achieved using the KT integrated with the SC (Figure 4 r-u). This configuration combines the advantages of predictive modeling with the strict linkage constraints of the SC to resolve a fully developed laminar profile. The resulting map demonstrates a dominant, high-velocity core that decays smoothly toward the vessel walls (Figure 4 r-u). By effectively eliminating extra-luminal noise and tracking artifacts seen in simpler methods, this integrated approach successfully reconstructs the theoretical parabolic profile required for accurate microvascular analysis.

**Figure 4:**
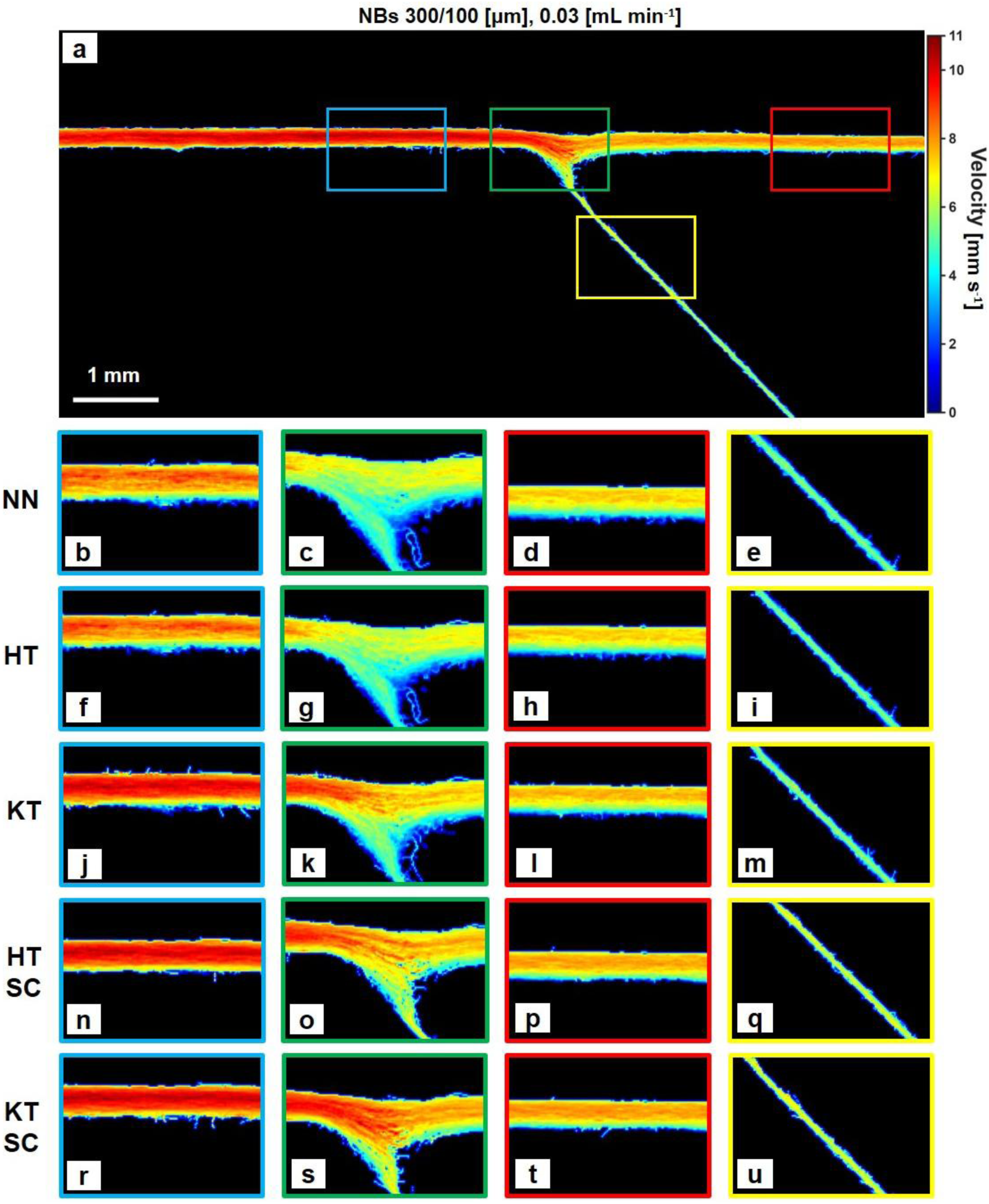
Comparative evaluation of ULM tracking algorithms. (**a**) Full velocity map of a gelatin phantom, reconstructed using KT SC; colored ROIs correspond to magnified regions in (b-u). **(b-e)** Nearest Neighbor (NN). **(f-i)** Hungarian Tracking (HT). **(j-m)** Kalman Tracking (KT). **(n-q)** Hungarian Tracking with smart cost matrix (HT SC). **(r-u)** Kalman Tracking with smart cost matrix (KT SC).

While the velocity maps provide a qualitative assessment of the reconstruction, a quantitative analysis is required to objectively measure the improvement in tracking fidelity. Five tracking configurations were compared to isolate the independent contributions of two algorithmic dimensions: the association strategy (greedy nearest-neighbor, globally optimal Hungarian assignment, and predictive Kalman filtering) and the cost function design (standard Euclidean distance versus the physics-informed SC). This factorial design enables attribution of performance gains to specific algorithmic choices rather than confounded improvements. To this end, statistical distributions of track morphology were computed for the entire dataset with all the tracks. For this analysis, the minimum track length filter was relaxed to 5 frames, compared to 15 in the visual maps. This relaxation was necessary to expose the raw performance of the tracking algorithms, including their tendency to generate short, fragmented, or erroneous tracks in high-density scenarios. A key biomarker for healthy microvascular flow in phantom channels is laminarity, which implies that trajectories should be smooth and direct. This characteristic is quantified by the Tortuosity Index (τ).

The baseline algorithms, NN and standard HT, exhibited a broad tortuosity distribution with a heavy tail extending up to τ = 8 (Figure 5 a,e). Crucially, the peak count at the ideal laminar value (τ ∼ 1) was relatively low, at approximately 1300 tracks for both the NN and the standard HT methods (Figure 5 a,e). When the data was filtered for tracks longer than 10 frames, the persistence of high-tortuosity outliers in these baseline methods remained evident, with a median tortuosity value of 1.4 (Figure 5 c,g). Significant improvements were observed when introducing the predictive motion models of the KT, which narrowed the distribution tail and increased the frequency of laminar tracks to approximately 3000 and 3500 counts for the raw and filtered datasets, respectively (Figure 5 i,k). The most substantial enhancement, however, occurred with the integration of the SC. By applying directional and brightness penalties to the linking cost, the SC configurations achieved a sharp peak at the ideal tortuosity value, effectively concentrating the distribution at τ ∼ 1 and reaching a peak count of 3000 tracks with HT SC and almost 4500 with KT SC, even before the data was filtered for tracks longer than 10 frames (Figure 5 m,q).

**Figure 5:**
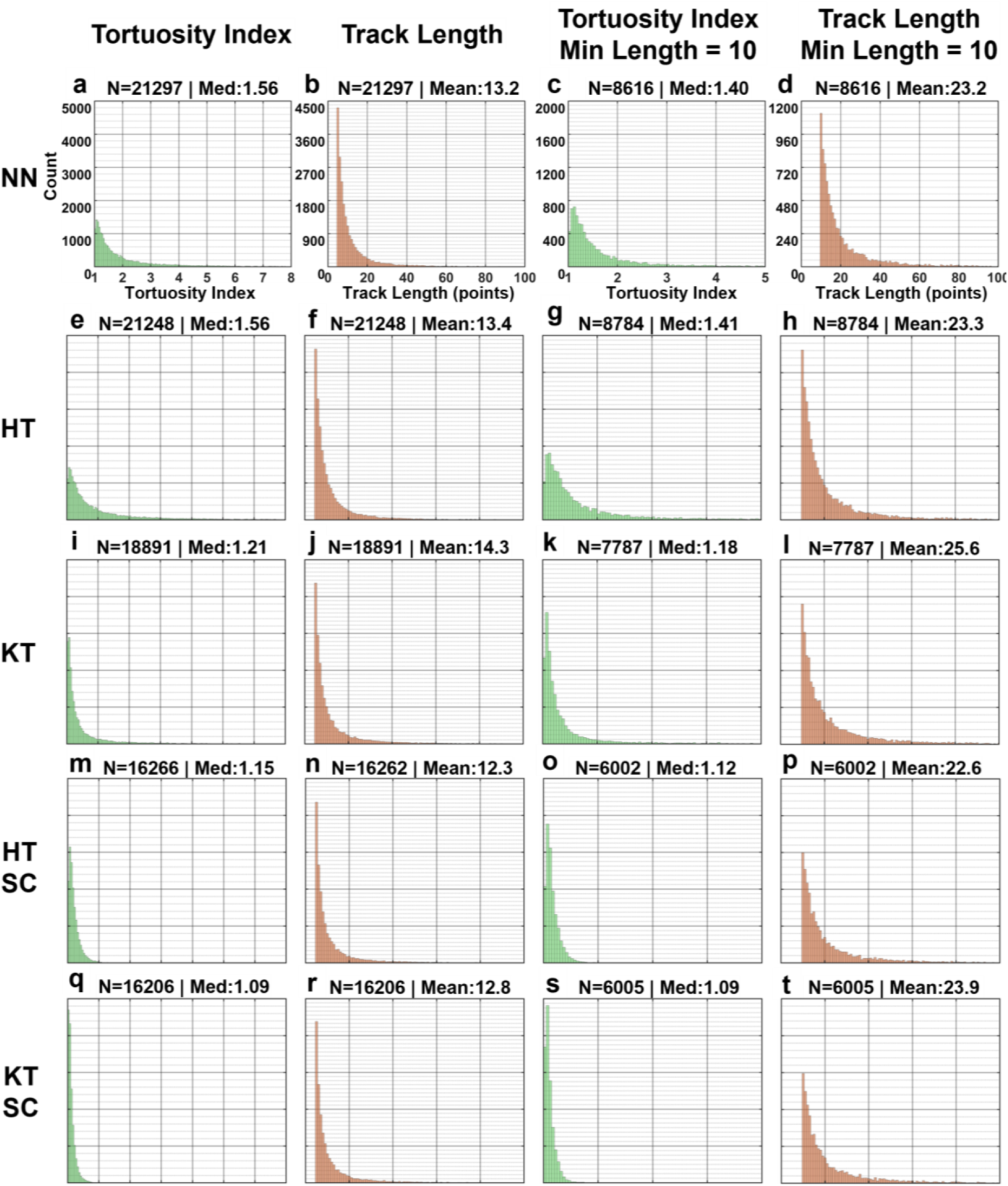
Statistical evaluation of tracking fidelity across algorithms. Histograms of Tortuosity (Columns 1, 3) and Track Length (Columns 2, 4) for five tracking configurations. **(a-d)** NN. **(e-h)** HT, **(i-l)** KT. **(m-p)** HT SC. **(q-t)** KT SC. Columns 1-2 show all tracks. Columns 3-4 show tracks filtered for length > 10 frames.

Furthermore, the total number of reconstructed tracks serves as a metric-based indicator for the algorithm’s selectivity versus its tendency toward fragmentation. The NN and standard HT methods produced the highest raw yields, with 21,297 and 21,248 tracks identified respectively (Figure 5 b,f). However, given the high tortuosity observed in these datasets, this surplus likely represents fragmented segments, where a single long vessel is erroneously broken into multiple short tracks and residual noise artifacts (Figure 5 d,h). The standard KT reduced this count to 18,891 tracks (Figure 5 j) with a median tortuosity of 1.21, while the inclusion of the SC resulted in the lowest overall yield of 16,206 tracks for the Kalman configuration (Figure 5 r). Despite the lower quantity, this configuration achieved the highest fidelity with the lowest overall tortuosity index of 1.09 for the unfiltered dataset (Figure 5 q), preserved after filtering for tracks > 10 frames (Figure 5 s). This reduction, combined with the increase in low-tortuosity tracks, confirms that the advanced algorithms successfully exchange quantity for quality. Finally, by filtering out spurious data and preventing the fragmentation of long vessels, the KT SC method reveals the physiological flow with the highest degree of confidence.

### Quantitative Hemodynamic Profiling and Velocity Validation

To validate the quantitative accuracy of NB as flow tracers, we performed a direct comparative analysis against MBs across three volumetric flow rates (0.01, 0.03, and 0.25 mL min^−1^). Evaluations were conducted in both symmetric (300-300 μm) and asymmetric (300-100 μm) bifurcation geometries cast within the gelatin phantoms. The velocity distribution analysis is performed across three key hydrodynamic zones, including the main channel prior to the bifurcation, the distal segment of the main channel, and the bifurcating branch (Figure 6).

**Figure 6:**
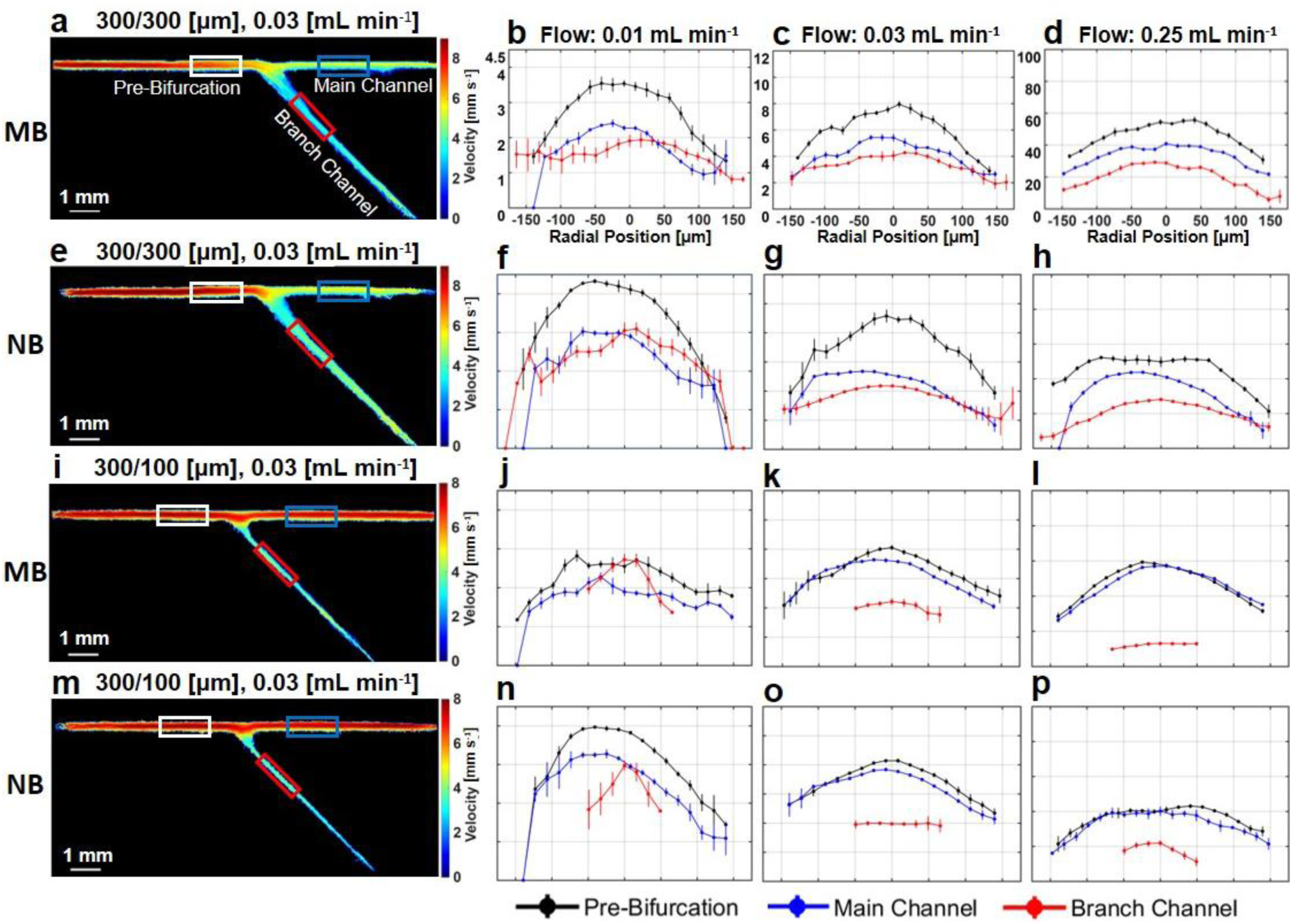
Quantitative velocity profiling of MB versus NB in symmetric and asymmetric bifurcation phantoms. (**rows a-h**) Comparative cross-sectional analysis across a symmetric 300-300 µm bifurcation and an **(rows i-p)** asymmetric 300-100 µm bifurcation, comparing MBs (rows a-d, i-l) to NBs (rows e-h, m-p). **(a, e, i, m)** Super-resolved velocity maps at 0.03 mL min^−1^. ROIs mark the pre-bifurcation (white), post-bifurcation (blue), and branch (red) segments. Cross-sectional velocity profiles are shown at **(b, f, j, n)** 0.01 mL min^−1^, **(c, g, k, o)** 0.03 mL min^−1^, **(d, h, l, p)** and 0.25 mL min^−1^. Axis labels and flow rate annotations appear only in the top row (b-d). Solid lines: measured ULM velocity profiles corresponding to the color-coded ROIs. Error bars represent ± SD across ROI axial positions per radial bin.

Initially, we evaluated the imaging performance using a symmetric bifurcation phantom 300-300 μm (Figure 6 a-h). Both contrast agents successfully reconstructed the expected laminar flow profiles. However, the agreement between the measured peak velocities in the pre-bifurcation segment (black profiles) and the calculated theoretical maxima varied as a function of the inlet flow rate. At the lowest flow rate (0.01 mL min^−1^), the measured velocities demonstrated high agreement with the theoretical peak velocity of 3.9 mm·s^−1^ (Figure 6 b,f). This correlation decreased slightly, yet remained comparable, at 0.03 mL min^−1^, where the theoretical peak is 11.7 mm·s^−1^ (Figure 6 c,g). At the highest tested flow rate (0.25 mL min^−1^), however, the measured peak velocities of approximately 60 mm·s^−1^ underestimated the theoretical maximum of 97.2 mm·s^−1^ by approximately 38%, despite an acquisition rate of 1000 Hz (Figure 6 d,h). A distinct hydrodynamic behavior was observed post-bifurcation. While the geometry is nominally symmetric, a velocity disparity emerged between the continuing main channel (Blue ROI) and the bifurcating branch (Red ROI). At lower flow rates (0.01 mL min^−1^), the velocities in both branches were comparable. However, at the highest flow rate of 0.25 mL min^−1^ (Figure 6 d,h), the velocity in the straight channel was notably higher than in the branching channel. This flow-rate-dependent velocity asymmetry in a nominally symmetric geometry was observed consistently for both contrast agents and is analyzed in the Discussion. Next, we examined the flow dynamics within an asymmetric bifurcation configuration 300-100 μm. The asymmetric phantom presented a high-resistance secondary channel (Figure 6 i-p). As expected, the velocity in the main channel post-bifurcation (Blue ROI) remained nearly identical to the pre-bifurcation velocity (Black ROI) across all flow rates. Conversely, the velocity in the 100 μm branch (Red ROI) was markedly lower, consistent with the preferential flow of fluid through the path of least resistance (the wide main channel). A minor reduction in the main channel velocity was observed downstream, corresponding to the small fraction of mass conservation diverted to the narrow branch.

### Comparative Analysis of Microvascular Partitioning and Flow Distribution

Having validated the hemodynamic fidelity of NBs through velocity profiling, the next phase of the analysis evaluated their ability to serve as accurate flow tracers within bifurcating microvascular geometries. Because NB and MB suspensions differ substantially in concentration, we employed the localization-based Partitioning Ratio metric to compare their flow distribution independent of contrast agent concentration. This approach isolates the physical transport behavior of the agents and enables direct comparison of their partitioning at vascular bifurcations. The results of this partitioning analysis, including super-resolved velocity maps for each geometry and pie charts quantifying the fractional distribution between branches, are presented across the various experimental regimes (Figure 7). Overall, the fractional distribution of NBs closely mirrored that of MBs across the majority of matched experiments (Figure 7 d-l vs. m-u), indicating that despite their substantially smaller size, NBs are subject to the same fluid dynamic forces that govern MB distribution in these channel diameters (100-500 μm). The sensitivity of the partitioning metric is confirmed by geometric comparison: symmetric phantoms yielded near-equilibrium distributions, while asymmetric geometries of 300-100 μm (Figure 7 g-i, p-r) and 500-200 μm (Figure 7 j-l, s-u) produced a marked shift for both agents toward the low-resistance branch, confirming that NBs faithfully trace the underlying hemodynamic resistance network.

**Figure 7:**
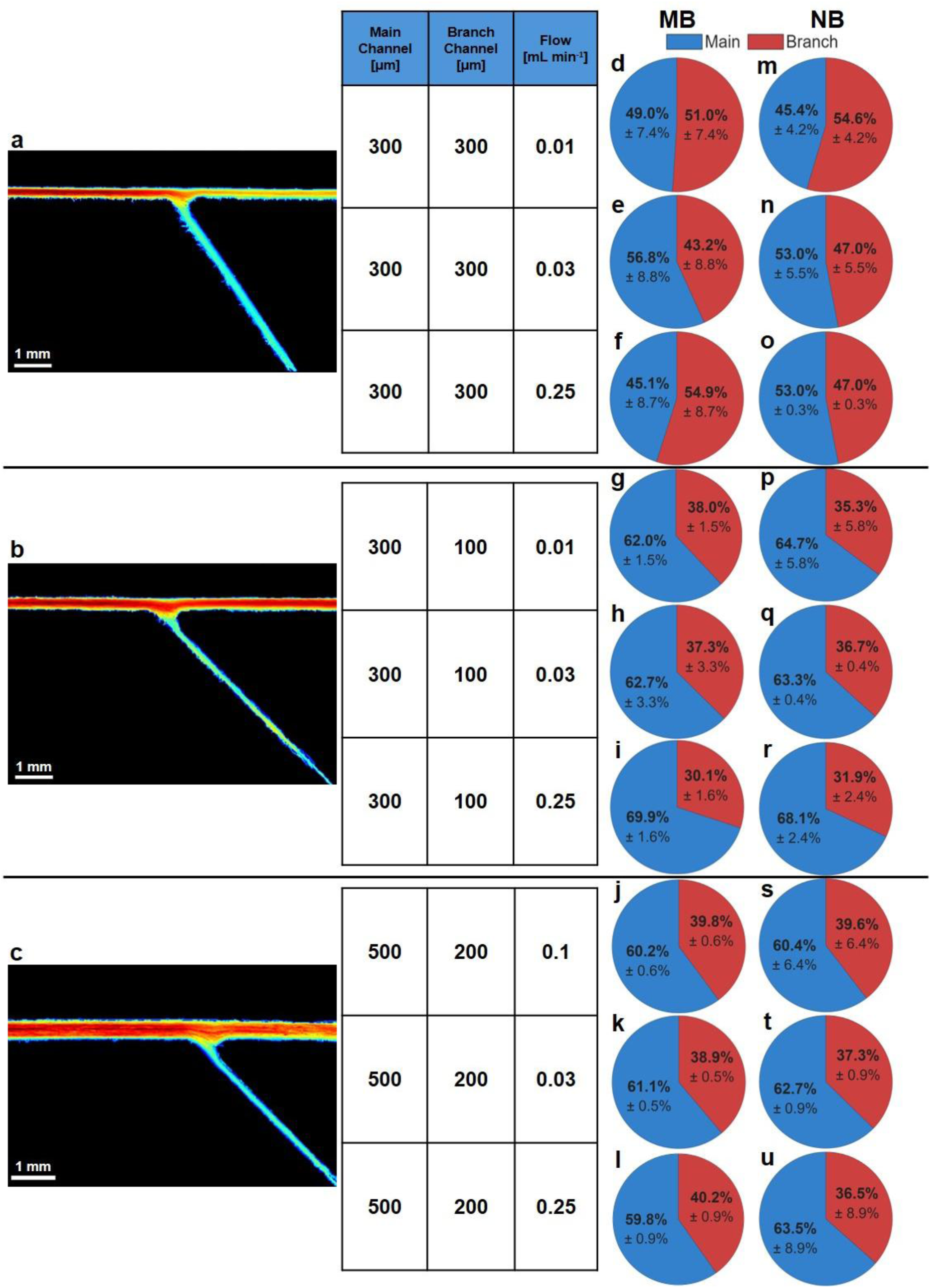
Comparative perfusion distribution of NB versus MB. (**a-c**) Super-resolved velocity maps for the three bifurcating phantom geometries. The central table lists the structural parameters and inlet flow rates for each row. **(d-l)** Pie charts showing the fractional distribution of MBs between the branch (red) and continuing main channel (blue). **(m-u)** Corresponding distributions for NBs. Values represent mean ± standard deviation (n = 3).

## Discussion

To the best of our knowledge, this study presents the first demonstration of ULM using sub-micron NBs as the primary contrast agent. Despite their substantially weaker acoustic scattering compared with conventional MBs [5], [16], NBs enabled accurate velocity reconstruction, flow partitioning measurements, and trajectory tracking across all tested phantom geometries. The close agreement between NB– and MB-based measurements indicates that NBs can serve as reliable flow tracers for ULM within the investigated vessel diameters (100-500 μm) Our results establish that NBs achieve hemodynamic mapping accuracy and vascular partitioning ratios almost identical to those of traditional MBs within 100-500 μm microvascular structures. The successful implementation of NB-based ULM highlights the importance of robust processing strategies when working with weakly scattering contrast agents [16]. Reliable localization and tracking of NB signals was facilitated by the ULM Master GUI, which provides an interactive environment for visualization and validation of the processing pipeline.

The comparison of tracking methodologies further demonstrated the importance of predictive motion models for NB-based ULM. While Nearest Neighbor and Hungarian tracking generated larger numbers of trajectories, these approaches produced shorter tracks and increased fragmentation [33], [41]. In contrast, the Kalman Tracker with Smart Cost Matrix (KT-SC) consistently yielded the lowest tortuosity values, approaching the ideal value of 1 observed for laminar flow. By incorporating motion prediction and trajectory-history information, KT-SC improved track continuity and reduced false associations, particularly under the high bubble densities required for NB imaging [35], [41]. These findings suggest that predictive tracking strategies can improve trajectory continuity in high-density localization datasets and may be beneficial for future NB-based ULM applications.

Several observations provide insight into the physical behavior of the phantom system itself. The systematic underestimation of peak velocity observed at the highest tested flow rate (0.25 mL·min⁻¹) is consistent with deformation of the gelatin channels under elevated internal pressure [23], [28]. Applying the continuity equation to the measured 38% velocity reduction predicts an effective increase in channel diameter from 300 μm to approximately 382 μm, corresponding to a 27% dilation. This estimate is consistent with the compliant nature of gelatin hydrogels and with the visible channel expansion observed during high-pressure flushing [28]. Similarly, the velocity asymmetry observed at the nominally symmetric bifurcation at high flow rates is consistent with flow redistribution at the junction, resulting in preferential flow through the straight branch [22], [23]. Importantly, both phenomena were observed consistently in the NB and MB datasets, supporting the conclusion that the measured differences originated from the physical flow environment rather than from contrast-agent-specific effects or tracking artifacts.

Several limitations should be considered when interpreting these results. First, the validation experiments were performed in vessel-mimicking channels ranging from 100 to 500 μm in diameter. While these dimensions are relevant to small vessels and bifurcating microvascular structures, they remain substantially larger than true capillary networks where differences arising from contrast-agent size may become more pronounced. Furthermore, the investigated vessel diameters were near or above the acoustic diffraction limit, such that the primary objective of this study was validation of NB localization and tracking rather than demonstration of a unique super-resolution capability enabled by NBs. Consequently, the present work establishes the feasibility and quantitative accuracy of NB-based ULM. Future studies will employ advanced fabrication techniques, including selective laser melting and powder-based 3D printing [28], [42], [43], to generate vessel geometries approaching true capillary dimensions [22], [44], enabling direct evaluation of size-dependent differences between MB and NB transport. Moreover, the present study was performed in blood vessel-mimicking phantoms. Translation to in vivo imaging will introduce additional challenges, including increased tissue clutter, attenuation, and physiological motion [6]. Future studies will therefore evaluate NB-based ULM in vivo and incorporate motion-compensation strategies to account for respiratory, cardiac, and tissue motion [34], [45]. In addition, the origin of individual localization events cannot be determined unequivocally. A detected signal may arise from a single NB or from multiple closely spaced NBs whose echoes are unresolved by the imaging system [46]. While the present study demonstrates reliable flow measurements despite this ambiguity, its impact on localization accuracy and density estimation remains to be systematically investigated. Finally, parameter optimization was performed interactively. Although this approach provides flexibility across diverse datasets and imaging conditions, it introduces operator dependence. Automated parameter-selection strategies and learning-based optimization approaches may reduce operator dependence and improve reproducibility across imaging platforms [47].

In terms of future directions, one particularly compelling application is the renal glomerulus, whose dense capillary architecture presents a longstanding challenge for conventional vascular imaging approaches [48]. Future studies will investigate whether NB-based ULM can improve visualization and quantification of glomerular microvascular structure and density, potentially enabling non-invasive assessment of renal pathologies such as focal segmental glomerulosclerosis [49], [50]. On the signal-processing side, future work will explore advanced beamforming and coherence-based imaging approaches, including Minimum Variance beamforming, Capon beamforming, and Short-Lag Spatial Coherence imaging, which may further improve bubble-to-clutter contrast and enhance NB detectability in low-SNR conditions [51]. Beyond lipid-shelled NBs, the framework developed here may also be applicable to emerging classes of weakly scattering ultrasound contrast agents. In particular, genetically encoded gas vesicles represent an intriguing opportunity for extending localization microscopy beyond vascular mapping toward molecularly targeted and genetically encoded ultrasound imaging [52], [53].

## Conclusions

This work demonstrates the feasibility of performing ULMith sub-micron NBs. Using custom wall-less gelatin flow phantoms, NB-based ULM achieved velocity reconstruction, flow partitioning, and tracking performance comparable to conventional microbubble-based ULM across vessel geometries ranging from 100 to 500 μm. To facilitate processing of low-SNR NB datasets, we developed the ULM Master GUI, an interactive framework that integrates the complete ULM processing pipeline and supports dataset-specific optimization. Together, these results establish NBs as reliable flow tracers for localization microscopy and provide an open-source framework for future investigations of NB-based ULM.

## Acknowledgements

This work was supported in part by the Israel Science Foundation under Grant 192/22, in part by an ERC StG under Grant 101041118 (NanoBubbleBrain), the Israel Cancer Research Fund (grant number 1286686), in part by the Alrov center for Digital Medicine, and in part by the Nicholas and Elizabeth Slezak Super Center for Cardiac Research and Biomedical Engineering at Tel Aviv University.

Generative AI tools (Claude, Anthropic and Gemini, Google) were used to assist with language editing and manuscript preparation. The authors reviewed and take full responsibility for all content.

## Author contributions

G.S. designed and performed the research, analyzed the data, and wrote the paper. Y.G. and M.B., assisted in experiments and data analysis. T.I. guided, advised, and designed the research and wrote the paper. All authors approved the final manuscript.

## Code Availability

The ULM Super-Resolution Toolbox, comprising the end-to-end processing pipeline (clutter filtering, sub-pixel localization, multi-algorithm tracking, and rendering) together with the ULM Master GUI optimization platform, is available at: https://github.com/grisha1998/ulm-super-resolution-toolbox. The toolbox is implemented in MATLAB and includes full documentation.

## Data availability

The datasets used in this study are available from the corresponding author on reasonable request.

## Table of Contents entry

Nanobubbles are established as a new class of contrast agents for ultrasound localization microscopy. Using an interactive processing framework and biomimetic wall-less microvascular phantoms, nanobubble-based imaging achieves accurate flow mapping and vascular partitioning, providing a foundation for super-resolution ultrasound imaging with nanoscale acoustic contrast agents.

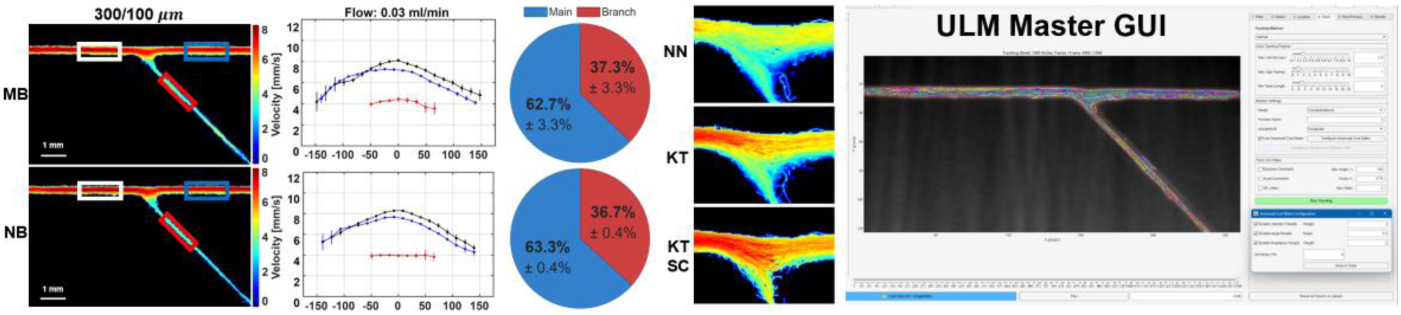

